# Minimal vertex model explains how the amnioserosa avoids fluidization during *Drosophila* dorsal closure

**DOI:** 10.1101/2023.12.20.572544

**Authors:** Indrajit Tah, Daniel Haertter, Janice M. Crawford, Daniel P. Kiehart, Christoph F. Schmidt, Andrea J. Liu

## Abstract

Dorsal closure is a process that occurs during embryogenesis of *Drosophila melanogaster*. During dorsal closure, the amnioserosa (AS), a one-cell thick epithelial tissue that fills the dorsal opening, shrinks as the lateral epidermis sheets converge and eventually merge. During this process, both shape index and aspect ratio of amnioserosa cells increase markedly. The standard 2-dimensional vertex model, which successfully describes tissue sheet mechanics in multiple contexts, would in this case predict that the tissue should fluidize via cell neighbor changes. Surprisingly, however, the amnioserosa remains an elastic solid with no such events. We here present a minimal extension to the vertex model that explains how the amnioserosa can achieve this unexpected behavior. We show that continuous shrinkage of the preferred cell perimeter and cell perimeter polydispersity lead to the retention of the solid state of the amnioserosa. Our model accurately captures measured cell shape and orientation changes and predicts non-monotonic junction tension that we confirm with laser ablation experiments.

**Significance Statement:** During embryogenesis, cells in tissues can undergo significant shape changes. Many epithelial tissues fluidize, i.e. cells exchange neighbors, when the average cell shape index increases above a threshold value, consistent with the standard vertex model. During dorsal closure in *Drosophila melanogaster*, however, the amnioserosa tissue remains solid even as the average cell shape index increases well above threshold. We introduce perimeter polydispersity and allow the preferred cell perimeters, usually held fixed in vertex models, to decrease linearly with time as seen experimentally. With these extensions to the standard vertex model, we capture experimental observations quantitatively. Our results demonstrate that vertex models can describe the behavior of the amnioserosa in dorsal closure by allowing normally fixed parameters to vary with time.

---

The developmental stage of dorsal closure in *Drosophila melangaster* occurs roughly midway through embryogenesis and provides a model for cell sheet morphogenesis (1–4). The amnioserosa (AS) consists of a single sheet of cells that fills a gap on the dorsal side of the embryo separating two lateral epidermal cell sheets. During closure, the AS shrinks in total area, driven by non-muscle myosin II acting on arrays of actin filaments in both the AS and actomyosin-rich cables in the leading edge of the lateral epidermis (5–8). Ultimately, the AS disappears altogether.

One might naively expect cells in the AS, which are glued to their neighbors by molecules such as E-cadherin, to maintain their neighbors, so that the tissue behaves like a soft, elastic solid even as it is strongly deformed by the forces driving dorsal closure. However, the time scale for making and breaking molecular bonds between cells (ms) is far faster than the time scale for dorsal closure (hours). As a result, cells can potentially slip past each other while maintaining overall tissue cohesion. Such neighbor changes could cause epithelial tissue to behave more as a viscous fluid on long time scales rather than an elastic solid, as it does during convergent extension (9). Vertex models (10–19) have provided a useful framework for how tissues can switch between solid and fluid behavior (14, 15), and have had remarkable success in describing experimental results (9, 20–23). These models make the central assumption that internal forces within a tissue are approximately balanced on time scales intermediate between ms and hours, and have successfully described phenomena such as pattern formation, cell dynamics, and cell movement during tissue development (24). Force balance is captured by minimizing an energy that depends on cell shapes. In the standard vertex model, a higher average cell shape index 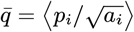 averaged over all cells, where *p*_*i*_ is the perimeter and *a*_*i*_ the area of cell *i*, corresponds to lower energy barriers and more fluid-like behavior, while a lower average shape index indicates more solid-like tissue properties. This framework links cell geometry, observable in imaging experiments, to collective tissue mechanics.

During dorsal closure, AS cells undergo significant shape changes. According to the standard vertex model, the observed high values of average cell shape index should render the tissue fluid (13–15). However, substantial experimental evidence indicates that the AS remains *solid* during dorsal closure with *no* neighbor exchanges (25–27). We have examined individual junction lengths using live embryo imaging in an extensive data set comprising 12 embryos, each with ∼100 cells and, in agreement with the literature, did not find any vanishing junctions, and hence any neighbor exchanges known as *T1 events*, except when cells left the AS (cell ingression). There are significant pulsatile contractions of the apical areas of AS cells (28). These fluctuations, as well as significant anisotropic strain due to the shrinking of the AS (see SI section M), should induce T1 events if the tissue were fluid. The fact that there are none implies that the tissue is a solid.

Vertex models might simply fail to describe tissue mechanics at this stage of development. The success of vertex models in describing many other tissues, however, begs the question: can the models be tweaked to capture tissue mechanics of the AS during dorsal closure, and could this point to an important physiological control mechanism? To address this question, we introduce a minimal extension to the standard vertex model that quantitatively captures cell geometry and tissue behavior from comprehensive experimental datasets obtained from timelapse microscopy recordings.

## 1. Modeling and experimental analysis

Our starting point is a standard two-dimensional cellular vertex model (10–19, 29, 30) (short introduction to vertex models in SI section A). The AS is represented as a single-layer sheet of polygonal cells that tile the entire area, as described below, with periodic boundary conditions. In our model, we approximate the shape of the AS tissue (Fig. 1A) with a rectangle whose long axis corresponds to the anterior-posterior axis of the embryo (Fig. 1B; for details see SI section B). During simulated dorsal closure, the positions of the vertices are continually adjusted to maintain the mechanical energy of the tissue at a minimum, or equivalently, to balance the forces exerted on each vertex. The mechanical energy of the standard vertex model is defined as

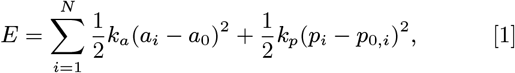

where *N* is the total number of cells, *p*_*i*_ and *a*_*i*_ are the actual cell perimeters and areas, *p*_0,*i*_ and *a*_0_ are the preferred cell perimeters and area, and *k*_*p*_ and *k*_*a*_ represent the perimeter and area elastic moduli of the cells, respectively. The first term penalizes apical area changes away from a preferred value, and can arise from cell height changes as well as active contractions in the medio-apical actin network at constant or near constant volume. The second term combines the effects of actomyosin cortex contractility with cell-cell adhesion, where *p*_0,*i*_ is the effective preferred cell perimeter (14). Without loss of generality, we chose *k*_*a*_ = *k*_*p*_ = 1 for all cells, as the area term acts merely as an energy offset, following the approach of Bi and Manning (14).

**Fig. 1.**
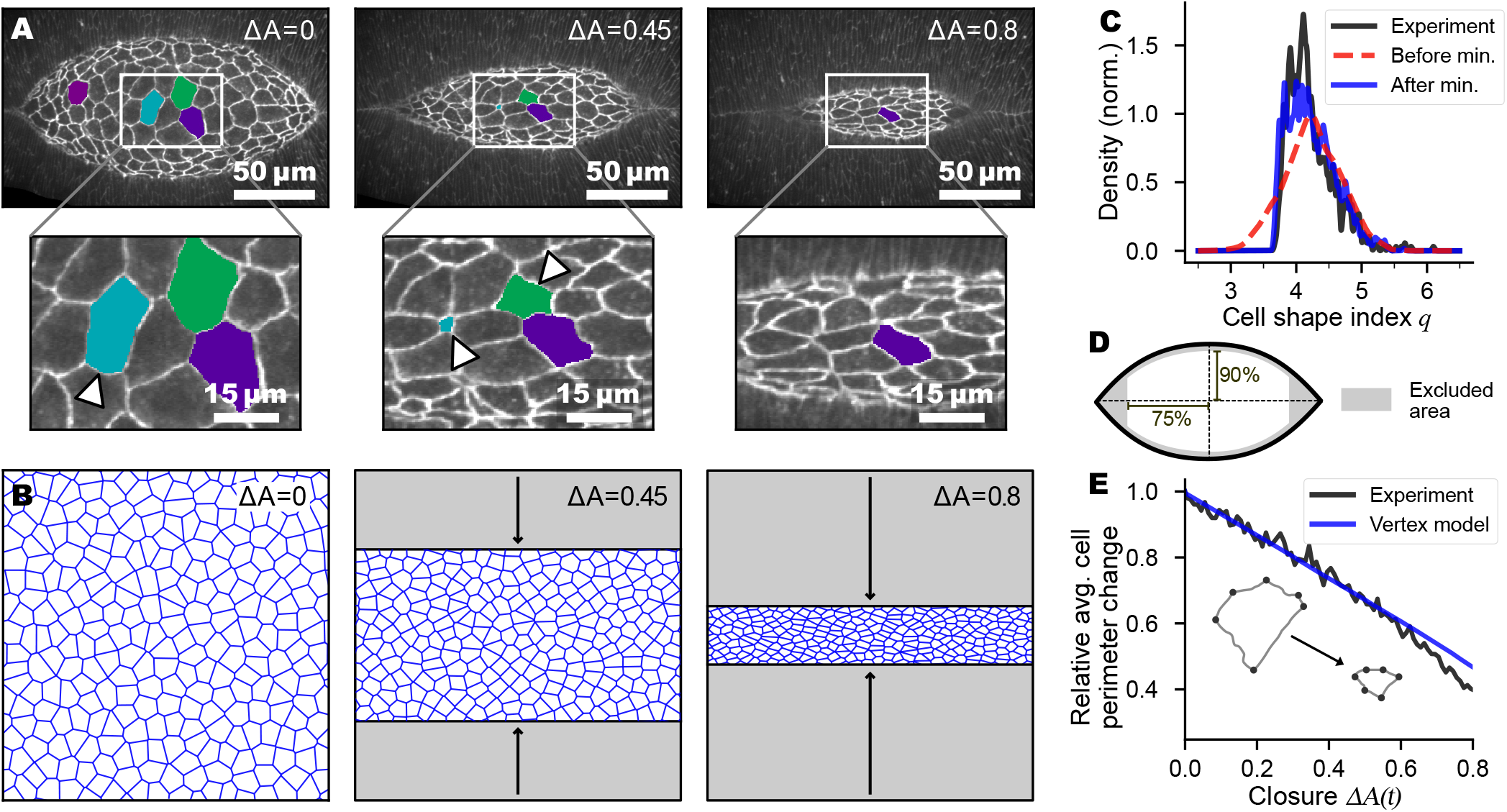
Experiment and vertex model for the amnioserosa during dorsal closure. (**A**) The geometry of the dorsal hole during early (left), middle (center), and late (right) dorsal closure. Enlargements show tissue with selected cells, several of which ingress (highlighted by triangles). (**B**) We model the dorsal closure process as a quasistatic uni-axial deformation. The geometry of the model is shown at the beginning (left), in the middle (center, at 45% closure), and towards the end (right, 80% closure) of the process. 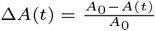 is the fractional change in total AS area of the closure process, where *A*_0_ is the AS area at the onset of dorsal closure. (**C**) An initial normal distribution of the preferred shape index of the model tissue (dashed red) with the standard deviation adjusted to be 0.45, leads to a distribution of the actual shape index after minimization (solid blue) that is in excellent agreement with the distribution of the experimentally observed shape index (solid black) at the beginning of dorsal closure. (**D**) Sketch of AS tissue regions included in model comparison (white center), with edge regions excluded (gray regions). (**E**) In the model, we reduce the preferred cell perimeter at a linear rate (blue) to capture the experimentally observed decrease of junction lengths (black). For comparison, we normalize the average perimeter by its value at the onset of the process. Inset: schematic representation of the reduction of cellular junction length and apical area during dorsal closure.

We used time-lapse confocal microscopy to image the entire dorsal closure process in E-cadherin-GFP embryos (Movie S1). We then used our custom cell segmentation and tracking algorithm to create time series of cell centroid position, area, perimeter, aspect ratio, and individual junction contour lengths for every cell in the AS.

During a substantial part of closure (Fig. 1A), the leading edges of the two flanking epithelial sheets approach the dorsal mid-line at a roughly constant rate (7), suggesting a uniform and anisotropic strain rate. Our experimental data confirmed this uniaxial strain pattern (SI section M). To mimic these dynamics, we linearly decreased the vertical height of the model tissue (Fig. 1B) as closure progresses. We enforce force balance, minimizing the mechanical energy after each deformation step of 0.125% of the initial height. Since closure rates varied from embryo to embryo, we measured progress during closure not in terms of *time*, but in terms of fractional change of the total area of the exposed AS (i.e. the dorsal opening), 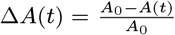, where *A*_0_ is a reference area of the AS early during closure. In many prior studies (31–33), the height of the AS has been used as a descriptor of closure progress. In Fig. S1 we demonstrate that both height and area of the AS decreased monotonically and approximately linearly with time, validating our use of Δ*A*(*t*) to mark the progression of closure. We began the analysis of each embryo at *A*_0_ = 11, 000 *μ*m^2^, a mid-stage of the full closure process at which the full AS was visible, so that we could average over multiple embryos. To exclude complex tissue boundary effects, we excluded cells at the AS borders and the regions at the canthi in the comparison between model and experiment (Fig. 1D, Movie S1).

AS cells reduce their perimeter (inset Fig. 1E) (34) during the closure process by removing a portion of junction material and membranes through endocytosis, while maintaining junction integrity (35, 36). The average perimeter shrinks at a constant rate in the experiments (Fig. 1E). We therefore assume in the model that the *preferred* perimeter *p*_0,*i*_ of each cell decreases linearly with Δ*A*(*t*) at the same rate (details see SI section B).

During dorsal closure, ∼ 10% of AS cells ingress into the interior of the embryo (3, 28, 37) (additional cells ingress at the canthi and adjacent to the lateral epidermis). In the model, we removed cells randomly at the experimentally measured rate so that roughly 10% of the AS cells disappeared over the course of dorsal closure.

At the onset of closure we find that cells in the AS exhibit considerable variability of the cell shape index 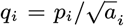 (Fig. 1C). In the model, we therefore introduce polydispersity in the cell shape index through a normal distribution of preferred cell perimeters *p*_0,*i*_ with standard deviation 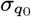. We hold 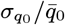 fixed throughout the process. In our minimal model, we further assume that the preferred cell area *a*_0_ is always the same for every cell. Thus we set *a*_0_(Δ*A*) = *A*_0_(1 − Δ*A*)/*N* (Δ*A*), where *A*_0_ is the total area at the onset of closure and *N* (Δ*A*) is the number of cells when the fractional area has decreased by Δ*A*. The distribution of actual shape index *q*_*i*_ after minimizing the mechanical energy in the model is in excellent agreement with the experiments (Fig. 1C).

Finally, we did not include cell motility in our simulations; this is justified since the amnioserosa is not attached to a solid substrate but is loosely attached through integrins to a flexible yolk (3). For more details on the model and experiments, see Materials and Methods and Supplementary Information.

## 2. Results

We tracked the following quantities during dorsal closure, in model and experiments: mean cell shape index 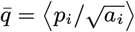 mean aspect ratio 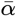 (see SI section D), orientational order parameter 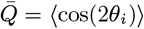 (38) (see SI section E) characterizing the degree of cellular alignment (where *θ*_*i*_ is the angle between the major axis of each cell *i* and the anterior-posterior axis, 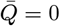 for randomly aligned cells and 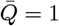 for cells perfectly aligned with the AP axis), standard deviation of cell shape index *σ*_*q*_ and standard deviation of the aspect ratio *σ*_*α*_.

We compare experimental data and simulations without any parameter modifications, adjustments, or rescaling with time. Considering the simplicity of the model, the agreement is remarkably good, both for cell shape index and cell shape index variability (Fig. 2A,B) as well as cellular alignment. The mean and standard deviation of cell aspect ratio agree equally well (Fig. 2C and SI Fig. S2). Our model slightly overestimates the orientation order of cells, possibly due to constraints on cells closer to the edge of the AS that we did not include in our simulations, despite excluding boundary cells from our analysis (Fig. 2D). As expected, the error bars (shaded region) in the experimental data, which represent variations between different embryos, are significantly larger than those in the simulation data, which represent only variations between initial configurations based on a single distribution of cell shape indices from the distribution measured over all embryos (Fig. 1C). In the experiments, there is intrinsic embryo-to-embryo variability that we did not include in our model for simplicity.

**Fig. 2.**
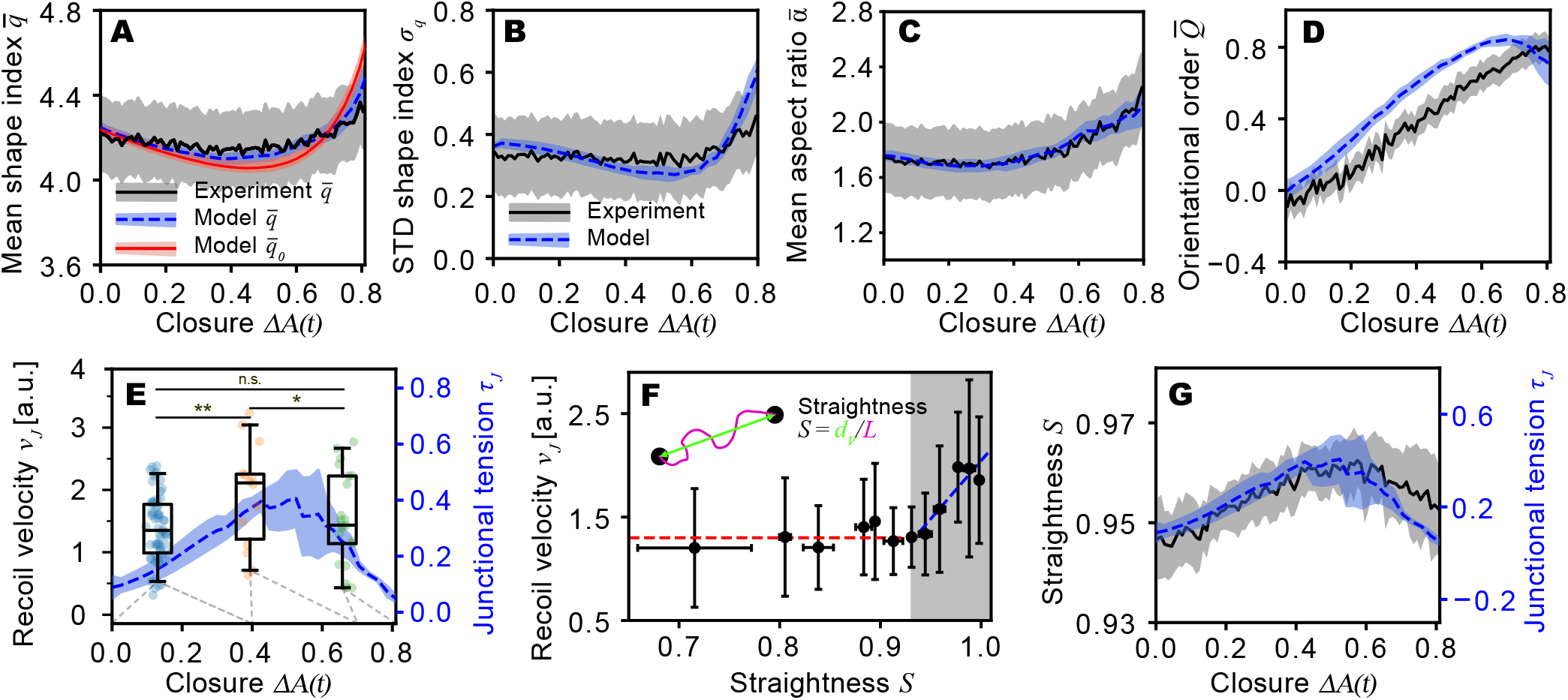
Results from experiment (black solid) and model (blue dashed). (**A**) A comparison of measured mean shape index 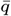 (blue curve) as a function of 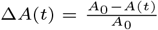. Here *A*_0_ is the AS area at the onset of dorsal closure and *A*(*t*) is the area as it shrinks during dorsal closure, so that Δ*A*(*t*) = 0 at onset. The red curve shows the model target shape index *q*_0_. The shaded regions indicate the standard deviation among 12 embryos (experiment) or 10 different initial configurations (model). (**B**) Comparison of cell to cell standard deviation of the shape index (*σ*_*q*_) during dorsal closure. (**C**) Comparison of cell aspect ratio during dorsal closure. (**D**) Orientational order parameter 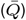 of the cells during dorsal closure. (**E**) Experimental initial junction recoil velocity (left y-axis) of the vertices after performing laser ablation of the junction, and predicted average cellular cortical tension 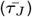(right y-axis) of the model during dorsal closure. The boxplots represent data across three intervals of Δ*A* (Δ*A* < 0.4, 0.4 ≤ Δ*A* < 0.7, Δ*A* ≥ 0.7). Whiskers extend to the 5th and 95th percentiles, while the boxes delineate the interquartile range, and the horizontal lines within the boxes indicate the median values. An ANOVA followed by a post-hoc Tukey’s HSD test was conducted to assess statistical significance (*: *p* < 0.1, **: *p* < 0.05). We performed and evaluated cuts of *N* = 97 junctions. (**F**) Average initial recoil velocity of vertices after laser cutting as a function of junction straightness (ratio of the inter-vertex distance (*d*_*v*_) to the junction length (*L*), see inset) immediately before cutting. Junction recoil velocity is independent of junction straightness (fitted with the red dashed line) until *S* = *d*_*v*_ */L* ≳ 0.93. The crossover point at *d*_*v*_ */L* ≈ 0.93 marks the intersection of the red and blue dashed lines; the latter fits the data points in the gray-shaded region, indicating that the recoil velocity increases strongly and approximately linearly with junction straightness in this regime. (**G**) Comparison of experimental junction straightness (left y-axis) and model cellular junction tension (right y-axis) during dorsal closure.

In experiment and model, the mean shape index 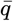 initially decreases, reaches a minimum at Δ*A* ≈ 0.55 and then increases (Fig. 2A). In the model, the behavior of 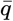 (blue curve) closely tracks the evolution of the preferred shape index, 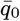 (red curve). Similar trends are seen in the width of the *q*-distribution, *σ*_*q*_ (Fig. 2B), which approximately tracks the mean 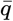 because of the constraint 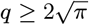, where the lower bound corresponds to a circle. The lower 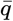, the more the distribution is squeezed against this lower bound, so the lower *σ*_*q*_. The increasing anisotropy during closure leads to greater alignment of cells along the anterior-posterior axis, reflected in an increased mean aspect ratio 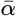 (Fig. 2C) and an increased orientational order 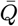 (Fig. 2D).

A strength of the vertex model is that it predicts not only cell shape and orientation distributions but also mechanical cell-level properties of the AS, including, for example, the average cell junction tension *τ*_*J*_, defined as (22, 39)

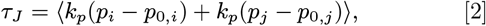

where *i, j* denote cells that share a given junction. Relative values of junction tension can be estimated experimentally from the initial recoil velocity *v*_*r*_ when a junction is severed using laser ablation (7, 12, 40–43). Our model predicts that the average junction tension *τ*_*J*_ rises until the fractional area change of the AS reaches Δ*A*(*t*) ≈ 0.55, and then decreases as dorsal closure continues (Fig. 2E). To test this prediction, we conducted laser cutting experiment at different stages of closure. We find that the recoil velocity changed in a non-monotonic manner (Fig. 2E), as the model predicts.

An alternative way to estimate junction tension from imaging data of unperturbed embryos is to analyze the straightness of junctions. A wiggly junction would be expected to be free of tension, whereas a straight junction should support tension. We define junction straightness as *S* = *d*_*v*_*/L* (Fig. 2F, inset), where *d*_*v*_ is the distance between vertices for a given junction and *L* is the contour length of the junction. We examined the relation between *S* and the initial recoil velocity upon cutting, *v*_*r*_, and observed that *v*_*r*_ is independent of *S* for *S* ≲ 0.93, but then rises linearly with increasing *S* above this threshold (Fig. 2E). It is reasonable to assume that junction straightness *S* is proportional to the tension *τ*_*J*_ predicted by our model. This is verified in Fig. 2G, showing the same non-monotonicity for both quantities with peaks occurring at Δ*A*(*t*) ≈ 0.55.

Why is the junction tension non-monotonic? In vertex models, junction tension and cell stiffness are related to cell shape index (14, 20, 21, 39). According to Eq. 2, junction tension is given by the difference between cell preferred perimeter *p*_0.*i*_ and the actual perimeter *p*_*i*_ for the two cells sharing a given junction. Below Δ*A* = 0.55, *p*_*i*_ − *p*_0,*i*_ increases with Δ*A*, leading to an increase of 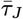. For Δ*A* ≥ 0.55, *p*_*i*_ − *p*_0.*i*_ decreases, leading to a decrease of 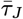.

A striking result of the standard vertex model (Eq. 1) is the prediction of a transition from solid to fluid behavior as the average shape index increases above 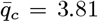 (14), in excellent agreement with a number of experiments in various epithelial tissue models (20, 21). Inspection of Fig. 2A shows that 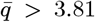 during the entire process of dorsal closure, suggesting that the AS should be fluid. However, the complete absence of T1 events (cell neighbor changes) strongly indicates that the AS is not fluid but solid.

For the tissue to behave as a solid, non-zero tension cell junctions must form continuous paths that extend across the entire system in all directions (44, 45)–in other words, they must *percolate*. The percolating network of tensed junctions creates a connected structure that can transmit forces across the tissue, thereby resisting deformation and conferring solid-like properties. Percolation requires the fraction of junctions with non-zero tension, *f*_*r*_ to be larger than a critical fraction *f*_*c*_. The rigidity transition can be driven either by altering *f*_*r*_ or *f*_*c*_, or both. Note that for random Voronoi tessellations in a square system, *f*_*c*_ ≈ 0.66 (46–48). The topology of such networks is similar to that of the standard vertex model (Eq. 1). We demonstrate that the percolation threshold *f*_*c*_ remains constant at ≈ 0.66 throughout dorsal closure, despite uniaxial deformation of the AS (Fig. 3A, SI section G). In both model and experiment the fraction of junction with non-zero tension remains above 0.7, showing conclusively that the AS remains rigid. Note that we have confirmed this result for our model by also counting T1 events (none), calculating the shear modulus and the deviatoric stress (SI sections L and N).

**Fig. 3.**
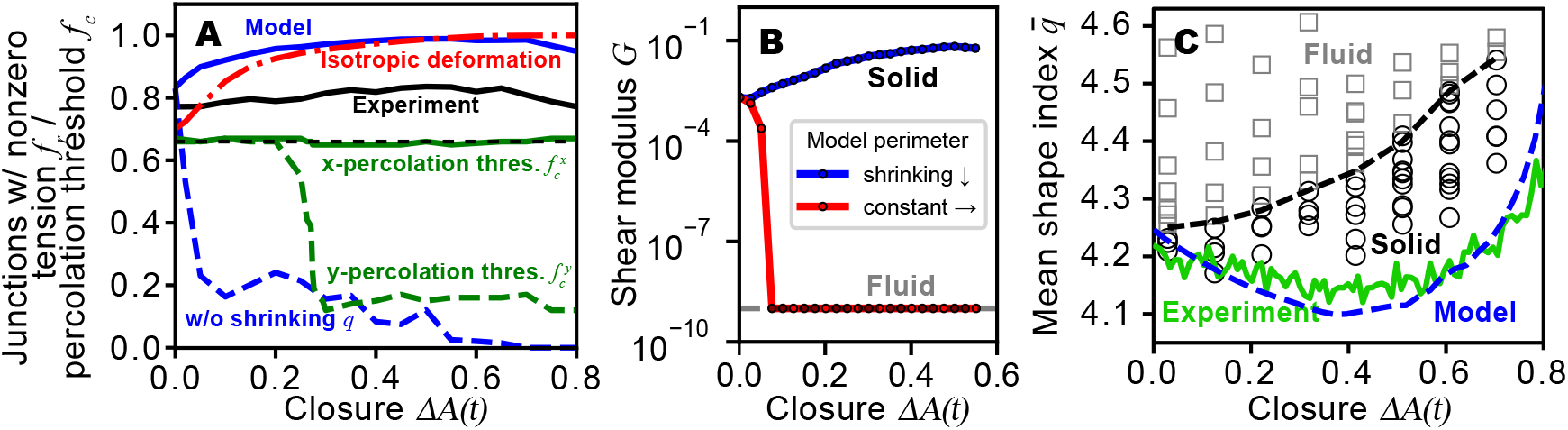
Percolation analysis, shear modulus, and phase diagram of AS during dorsal closure. (A) Percolation analysis of tissue rigidity, showing the fraction of junctions with non-zero tension in the model (blue line) and experiment (black line). Additional lines represent isotropic deformation (red dashed), model without shrinking preferred cell perimeters (blue dashed), and percolation thresholds in x-direction (green solid) and y-direction (green dashed). (B) Shear modulus of the minimal model throughout dorsal closure. The blue line shows that the shear modulus is nonzero for the model with shrinking preferred cell perimeter, while the red line shows that the system fluidizes (the shear modulus vanishes) when the preferred perimeters are held constant. (C) Phase diagram in 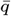 vs. Δ*A* space, illustrating solid (open black circles) and fluid (open gray squares) states. The solid-fluid transition is denoted by the black dashed line. Trajectories of both the experiment (green dashed line) and model (blue dashed line) remain within the solid phase throughout dorsal closure.

Fig. 3C summarizes our results in the form of a phase diagram of our model, obtained by evaluating the fraction of rigid junctions *f*_*r*_ at discrete values of mean cell shape index and ΔA (details on generation of phase diagram see SI section J). The phase boundary (black dashed curve) corresponds to the percolation transition of nonzero-tension junctions, *f*_*r*_ = *f*_*c*_ ≈ 0.66.

The system (blue dashed curve) always remains in the solid phase, below the black dashed line, consistent with the experimental observations.

Why does 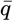 for the system (blue dashed line) initially decrease and then later increase with increasing Δ*A*(*t*)? This trend, shown also in Fig. 2A, is driven by the behavior of the preferred shape index (red curve in Fig. 2A). Minimization of the energy tends to drag 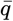 towards its preferred value. We can then ask why 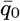 is non-monotonic in Δ*A*(*t*). Recall that the shape index for a cell is 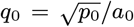, where *p*_0_ is the preferred perimeter of the cell and *a*_0_ is the preferred area. During closure, *p*_0_ shrinks linearly with increasing Δ*A*(*t*) as shown in Fig. 1E. However, *a*_0_ = *A*(*t*)*/N* (*t*), where *A*(*t*) is the area of the system and *N* is the number of cells at time *t*, also shrinks with increasing Δ*A*(*t*). The net result is the red curve shown in Fig. 2A.

We now turn to the central question of this paper–why does the system remain in the rigid state throughout closure? To understand this, we must address the behavior of the phase boundary 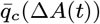 (black solid line in Fig. 3C). Note that the standard vertex model (Eq. 1) would predict simply 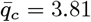. In our model, even at the beginning of the process, the phase boundary lies at 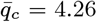. This differs from the standard value of 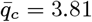 because the preferred perimeters are heterogeneous (44) with a distribution given by the red dashed curve in Fig. 1C. However, we hold the relative width of the preferred perimeter distribution fixed during the process and the shape index of the system eventually grows far above 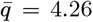, yet the system remains in the solid phase. The reason why the tissue *remains* solid during closure is therefore not the heterogeneity in preferred cell perimeters.

A second possibility is that during dorsal closure, orientational ordering of cells along the anterior-posterior axis further shifts the phase boundary upwards (9) (for a detailed analysis of orientational alignment see SI section H). However, the maintenance of the solid phase in our model does not stem primarily from orientational ordering. We show this by performing isotropic compression instead of uniaxial compression to shrink the area of the system (SI section I). If the system is subjected to isotropic compression, no orientational order is induced (Fig. S5B), as one might expect. Nevertheless, we find that our model predicts solid behavior even for isotropic deformation, showing that uniaxial deformation is not needed for this aspect. We note that although uniaxial compression is not important for maintaining tissue in the solid phase during closure, it is needed to obtain the trends in Fig. 2A-D,G, since trends in 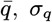 and junction tension *τ*_*J*_ with closure fail to agree with experimental results if we apply isotropic deformation (Fig. S5A,D,C).

Yet another factor in our model that could affect the location of the solid-fluid phase boundary is cell ingression. However, we find that cell ingression at the levels seen experimentally (<10%,(28, 49)) has almost no effect on the location of the solid-fluid transition in the model.

The reason why the phase boundary shifts upwards in 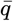 is that the preferred cell perimeter shrinks with increasing Δ*A*(*t*) as in Fig. 1E. We demonstrate this unambiguously by keeping the preferred cell perimeter fixed. In that case, we find that the tissue fluidizes rapidly (blue dashed line in Fig. S3B) as the total area is decreased, with ∼ 2,000 T1 events observed as Δ*A*(*t*) increases from 0 to 0.8. We further confirm this finding by evaluating the shear modulus of our model (Fig. 3B), which drops rapidly to zero with increasing Δ*A* when the preferred perimeters are held constant (red curve).

Finally, one might ask whether asynchronous periodic fluctuations of preferred AS cell areas by ∼ 10%, as observed experimentally during closure, which have been neglected in the minimal model, might affect the phase boundary. We have checked this and find that such fluctuations lead only to a weak softening effect, not fluidization (SI section K).

## 3. Discussion

We find experimentally that the AS exhibits no cell neighbor exchanges, or T1 events during dorsal closure. The absence of T1 events does not in itself necessarily imply the tissue is solid. If there is no anisotropic strain applied and there are no fluctuations of cell vertices, due either to temperature or active forcing in the form of myosin-driven contractions or cell propulsion, then there would be no T1 events even in the fluid phase. Conversely, the mere presence of T1 events does not imply a fluid state. A solid phase under sufficient anisotropic strain could undergo plastic flow via T1 events.

In this case, we have demonstrated that there is significant anisotropic strain in the AS (SI section M). The AS is also subjected to strong pulsatile contractions during closure (28).

If the AS were fluid or even a sufficiently weak solid, these would drive T1 events. The fact that there are no T1 events is therefore convincing evidence that the AS is solid.

In the AS, cells exhibit highly elongated shapes (Fig. 2A,C). According to the standard vertex model, such highly elongated cells should facilitate low energy barriers for neighbor switching, theoretically resulting in fluid-like tissue behavior at the observed mean cell shape index 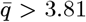 (9, 15, 20, 21, 50). Notably, this fluidization at elongated cell shapes has also been observed in other models, e.g., spring-edge models (51). However, this prediction is in remarkable contradiction with the experimentally observed tissue behavior.

Our minimal model, incorporating parameters based on experimental results, such as cell shape heterogeneity, shrinkage of preferred cell perimeters, and uniaxial tissue constriction, not only predicts the tissue’s solid phase but also faithfully reproduces a wide range of features of an extensive set of experimental dorsal closure data: cell shape and orientational order, and junction tension, which we inferred passively from image data due to the linear relationship between junction straightness and initial recoil velocity in laser cutting experiments.

We find that cell shape polydispersity and active shrinking of the preferred cell perimeters are the two critical factors that enable the tissue to remain solid in spite of extensive cellular and tissue shape changes. The effect of the shrinking of preferred perimeters is particularly important because it shifts the solid-fluid phase boundary from its initial value of 4.26 at the onset of closure to a far higher value of 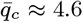 during closure. These results imply that the solid character of the AS originates from active processes that regulate cell perimeter, including junction complexes and the components of the cell cortex.

This finding raises two questions for future research. First, *how* is the removal of junction material specifically regulated in cells? Second, *why* might it be important for the AS to remain in a solid phase? Perhaps solid behavior during dorsal closure is simply a holdover from the preceding developmental stage of germ band retraction (52). Laser ablation experiments (53) suggest that the AS plays an important assistive role in uncurling of the germ band by exerting anisotropic tension on it. Such anisotropic stress requires the AS to be solid, not fluid. An interesting future direction for experimental and vertex model studies is to establish whether the AS is solid throughout germ band retraction as well as dorsal closure.

Our results show that vertex models are more broadly applicable than previously thought. Despite the many complex active processes that occur during dorsal closure, we find that only one of them–the active shrinking of a normally-fixed parameter, namely the preferred perimeter–is needed in order to quantitatively describe our experimental observations. Similar variation of normally-constant parameters has been shown to allow other systems to develop complex responses not ordinarily observed in passive non-living systems. These include negative Poisson ratios (54, 55) and allostery (56) in mechanical networks, greatly enhanced stability in particle packings (57), and the ability to classify data and perform linear regression in mechanical and flow networks (58) as well as laboratory electrical networks (59). More generally, the mechanical behavior of epithelial tissues during development is extraordinary when viewed through the lens of ordinary passive materials. It remains to be seen how much of that behavior can be understood using “adaptive vertex models” (60) within a framework that replaces ordinarily fixed physical parameters such as the preferred perimeter with adaptive degrees of freedom that vary with time.

## Materials and Methods

Flies were maintained using standard methods, and embryos were collected and prepared for imaging and laser surgery as previously described (61–63). Cell junctions were labeled via ubiquitous expression of DE-cadherin-GFP (64). Images were captured using Micro-Manager 2.0 software to operate a Zeiss Axiovert 200 M microscope outfitted with a Yokogawa CSU-W1 spinning disk confocal head, a Hamamatsu Orca Fusion BT camera, and a Zeiss 40X LD LCI PlanApochromat 1.2 NA multi-immersion objective (glycerin). Due to the embryo’s curvature, multiple z planes were imaged for each embryo at each time point to observe the dorsal opening. We recorded stacks with eight z-slices with 1 μm step size every 15 s throughout the closure duration, with a 100 ms exposure per slice.

Two-dimensional projections of the AS tissue were created from 3D stacks using DeepProjection (65). The height difference along the Z-axis between the center and the edge of the amnioserosa is less than 3 μm, which is negligible compared to the total tissue width of approximately 80 μm in the XY plane. Consequently, distortions of AS cell shapes due to the 2D projection are minimal. A custom Python algorithm was used to segment and track individual AS cells throughout dorsal closure: Briefly, binary masks of the AS cell boundaries and the AS tissue boundary (leading edge) were first predicted from microscopy movies using deep learning trained with expert-annotated dorsal closure specific data (66). Second, individual AS cells were segmented and tracked throughout the process using the watershed segmentation algorithm with propagated segmentation seeds from previous frames. Finally, for each cell, area, perimeter, shape index, aspect ratio and orientation in relation to the AS anterior-posterior axis were quantified over time. Based on the binary mask of the leading edge, we segmented the dorsal hole/AS shape, fitted an ellipse to it at each time point, and located the centroid position of each cell with respect to the long and short axis of the ellipse. This allowed us to precisely identify cells in the amnioserosa center (within 75% of the semi-major axis and 90% of the semi-minor axis), and exclude peripheral cells from comparisons between model and experiment. The straightness *S* of cell-cell junctions was quantified by segmenting the contour and end-to-end lengths of individual junctions using a custom graph-based algorithm. Laser surgery was performed on a Zeiss Axio Imager M2m microscope equipped with a Yokogawa CSU-10 spinning disk confocal head, a Hamamatsu EM-CCD camera and a Zeiss 40X, 1.2 NA water immersion objective. Micro-Manager 1.4.22 software controlled the microscope, the Nd:YAG UV laser minilite II (Continuum, 355 nm, 4 mJ, 1.0 MW peak power, 3–5 ns pulse duration, 10 Hz, (62)) and a steering mirror for laser incisions. In each embryo (N = 48), 1 to 2 cuts of approx. 5 μm length with a laser setting at 1.4 μJ were performed in the bulk of the AS at different stages of closure (62, 67, 68) (Fig. S6A,B). The response of the AS was recorded prior to (∼ 20 frames), during (∼ 4 frames) and after (∼ 576 frames) the cut at a frame rate of 5 Hz. The junction straightness *S* of each cut junction was quantified prior to the cut by manually tracing junction end-to-end length and junction contour length using ImageJ. Then, to analyze the initial recoil velocity, the motion of the vertices adjacent to the cut junction was followed in a kymograph perpendicular to the cut (line thickness 2 μm, Fig. S6A-C). On the basis of the kymograph, the distance *d*(*t*) between the two vertices of the severed junction was quantified manually over time using ImageJ. A double exponential function *a*_0_ exp(*b*_0_*t*) + *c*_0_ exp(*d*_0_*t*) + *e*_0_ was fitted to *d*(*t*) (Fig. S6D). The initial slope of this function at *t* ∼ 0 corresponds to the initial recoil velocity *v*_*r*_.

For the vertex model, we used the open-source CellGPU code (69). Analysis and illustration of model and experiment data was performed with custom Python scripts. Simulation code is available at https://github.com/indrajittah/Vertex-Model-Amnioserosa-Dorsal-Closure. Data associated with this study are available upon request.

## ACKNOWLEDGMENTS

We thank M. L. Manning, Sadjad Arzash, and S. R. Nagel for instructive discussions. This project was supported by NIH through Awards R35GM127059 (DPK) and 1-U01-CA-254886-01 (IT), NSF-DMR-MT-2005749 (IT, AJL) and by the Simons Foundation through Investigator Award #327939 (AJL). AJL thanks CCB at the Flatiron Institute, as well as the Isaac Newton Institute for Mathematical Sciences under the program “New Statistical Physics in Living Matter” (EPSRC grant EP/R014601/1), for support and hospitality while a portion of this research was carried out.

