## Supplementary file for "Minimal vertex model explains how the amnioserosa avoids fluidization during *Drosophila* dorsal closure"

**This PDF file includes:**

Figs. S1 to S9
SI References

**A. Short introduction to vertex models.** Vertex models for tissues were motivated by two-dimensional models of dry foams made up of polygonal bubbles. In such foams, the vertices of bubbles tend to adjust their positions to balance forces arising from line tensions at the boundaries between adjacent bubbles; this force balance is achieved by minimizing the energy cost associated with bubble-bubble boundaries (1, 2). Tissue imaging typically provides time-evolving 2D projections of the tissue layers. 2D modeling of 2D projections in vertex models is appropriate, since tension is provided by a 1D ring of acto-myosin, and adhesion is provided by E-cadherin also in a ring-like geometry. One can visualize such a model as a tightly packed layer of elastic polygons, having a finite relaxed area, with a line tension around the circumference and "slippery" adhesiveness to their neighbors that prevents gaps, but allows sliding. With these assumptions, computer simulations show that barriers preventing cells from sliding past each other are low when cells are elongated, i.e., the tissue reacts like a fluid to external forces on time scales determined by the remaining friction. Barriers are high, in contrast, when cells are more rounded, i.e., the tissue reacts like a solid under external forces in the sense that the tissue may deform, but cells maintain their neighbors. This type of very simple modeling has been extraordinarily successful in providing quantitative predictions of the decrease and loss of solid character in a wide range of epithelial tissues (3–6) and monolayer cell cultures. It was found that one parameter, mean cell shape index ($\bar{q} = \langle p_i/\sqrt{a}_i \rangle$ ), where $p$ is the cell perimeter and $a$ is its area, was dominant in determining phase state in many systems (3–5). Within the standard vertex model, the AS is a two-dimensional plane comprising irregular polygons that represent cells and that display no overlaps or gaps, as shown in Fig. 1A,B in the main text. The positions of the vertices adjust to minimize the mechanical energy

$$E = \sum_{i=1}^{N} \frac{1}{2} k_a (a_i - a_0)^2 + \frac{1}{2} k_p (p_i - p_{0,i})^2. \qquad [1]$$

Here $N$ is the total number of cells, $p_i$ and $a_i$ are the actual perimeter and area of cell $i$ and $p_{0,i}$ and $a_0$ are its preferred perimeter and area. $k_a$ and $k_p$ represent the area and perimeter moduli of the cells. The first term in Eq. 1 represents the area elasticity of the cells. According to this term, there are restoring forces if the area of cell $i$, $a_i$, differs from its preferred area, $a_0$. We note that there are medioapical arrays of actomyosin that condense and relax as pulsing AS cells contract and expand (7). In the vertex model, the medioapical arrays would control the preferred area, $a_{0,i}$. In our model, we neglect such effects, taking $a_{0,i}$ to be constant in time and the same for every cell ($a_{0,i} = a_0 = \text{const}$).

The second term in Eq. 1, involving the cell perimeter, originates from the sum of two contributions

$$k_{p,i} p_i^2 + \gamma_i p_i = \frac{1}{2} k_p (p_i - p_{0,i})^2 \qquad [2]$$

where $p_{0,i} = -\gamma_i (2k_{p,i})$ is the effective target cell perimeter (8). The first term on the left side of Equ. 2 approximates active contractility of the actomyosin sub-cellular cortex. According to this term, the cortex acts like a spring that prefers each cell to have a perimeter of zero (complete contraction); this contributes a term proportional to the square of the perimeter (9). The second contribution (second term on left side of Eq. 2) arises from cell-cell adhesion and cortical tension which are proportional to the perimeter. They combine to give an effective line tension $\gamma_i$ that penalizes a nonzero perimeter if $\gamma_i > 0$ (if contractility dominates) or rewards a nonzero perimeter if $\gamma_i < 0$ (if cell-cell adhesion dominates) (8–10).

**B. Details of the amnioserosa vertex model.** The initial AS cell sheet is modeled as a two-dimensional rectangular plane sheet with box size $L_x \times L_{y,\text{initial}}$ that is tiled with $N_{\text{initial}} = 256$ cells with no gaps or overlaps. We use periodic boundary conditions throughout the process. For simplicity, we set the preferred area for each cell $i$ to be the same for all cells: $a_{0,i} = 1$ .

The experimentally observed heterogeneity in cell shape index of the AS tissue is introduced in our model through the preferred perimeters of cells ($p_{0,i}$ for cell $i$). The initial values of $p_{0,i}$ are drawn from a Gaussian distribution with mean $\mu = 4.24$ and standard deviation $\sigma = 0.45$. We then minimize the energy using the FIRE algorithm (11) until the residual force on each vertex drops below $10^{-6}$ to obtain the initial state of the tissue. After minimization, we compare the distribution of shape index from both simulation and experiment (Fig. 1C, main text).

We approximate the dorsal closure process by applying uni-axial quasi-static compression to our model tissue. We hold $L_x$ fixed and with each step, we decrease $L_y$ by $\epsilon = 0.01$ and re-equilibrate the system (minimize the mechanical energy) so that there is force balance at every vertex.

During the process, we also decrease all of the preferred cell perimeters with a linear rate corresponding to the average perimeter drop observed in the experiments ($m_p = 2.76$) (Fig. 1E, main text):

$$p_{0,i}(\Delta A(t)) = p_{0,i}(\Delta A(0)) - m_p \Delta A(t). \qquad [3]$$

**C. Relation between AS height and AS area.** In Fig. S1 relation between the AS height and area is demonstrated.

**D. Cell shape index and aspect ratio.** The shape index of cell $i$ is defined as $q_i \equiv p_i/\sqrt{a_i}$, where $p_i$ is the cell perimeter and $a_i$ is the cell area. The mean cell shape index is $(\bar{q}) = \langle \frac{p_i}{\sqrt{a_i}} \rangle = \frac{1}{N} \sum_{i=1}^{N} \frac{p_i}{\sqrt{a_i}}$.

The aspect ratio of a cell is defined as

$$\alpha = \frac{\sqrt{\frac{1}{\lambda_1}}}{\sqrt{\frac{1}{\lambda_2}}} \qquad [4]$$

where $\lambda_1$ and $\lambda_2$ are smallest and largest eigenvalues of the shape tensor $I$, defined as

$$I = \begin{pmatrix} I_{XX} & I_{XY} \\ I_{XY} & I_{YY} \end{pmatrix} \qquad [5]$$

Here, $I_{XX} = \iint_R y^2 dA$, $I_{YY} = \iint_R x^2 dA$, and $I_{XY} = \iint_R xy dA$. The tensor $I$, calculated as in Ref. (4), corresponds to the moment-of-inertia tensor in the case where the mass density per unit area is constant. Thus $I$ measures the second area moment of the polygon representing the apical surface of the cell, relative to a fixed point, weighting each infinitesimal area element of the cell equally. In this case, the fixed point is the center of the cell, as defined by the first moment (the mean) of the area distribution of the polygon. Thus, the second moment measures the variance of the area distribution. In 2-dimensions, the tensor is a $2 \times 2$ matrix as defined above and its two eigenvalues $\lambda_1$ and $\lambda_2$ along two directions that in the case of an ellipse would correspond to the minor and major axes, respectively.

**E. Cell orientational order parameter.** During dorsal closure, cells tend to align along the anterior-posterior axis. According to the standard vertex model, this alignment should enhance the solid character of the tissue (5). To better understand the relationship of solidity to the collective alignment among aminoserosa cells, we quantify their degree of orientational order (12–14). This tells us the degree to which cells align relative to the anterior-posterior axis. The 2D orientational order parameter $\bar{Q}$ is (15)

$$\bar{Q} = \frac{1}{N}\sum_{i=1}^{N}\cos(2\theta_i) \qquad [6]$$

where $\theta$ is the angle of the major axis of the cell relative to the anterior-posterior axis. As described in Sec D, the direction of the cell's major axis is given by the largest eigenvector of the shape tensor $I$. Note $\bar{Q} = 0$ when cells are randomly aligned in the tissue (the isotropic case) and $\bar{Q} = 1$ if all cells are perfectly aligned along the anterior-posterior axis.

**F. Relation between mean and standard deviation of cell aspect ratio.** Similar to the cell shape index $q$, the cellular aspect ratio $\alpha$ also characterizes cell shape (4). Fig. S2 demonstrates that our model successfully captures experimental observations for both the mean and standard deviations of the tissue aspect ratio.

**G. Rigidity percolation in uniaxially deformed tissue.** Uniaxial deformation during dorsal closure produces anisotropy in the tissue which can affect the percolation threshold. To probe how the percolation threshold $f_c$ depends on the uniaxial deformation, we take the model tissue at each value of $\Delta A(t)$ and randomly assign each cell-cell junction to have nonzero tension with probability $f$ (Fig. S3A). Percolation of the system as a whole requires percolation in both the x and y directions. Therefore, the percolation threshold of the system is given by $f_c = \max[f_c^x, f_c^y]$ (Fig. S3C,D). To determine the percolation thresholds $f_c^x$ and $f_c^y$ at different stages of closure $\Delta A$, we assessed the probability of obtaining a system-spanning connected path of non-zero tension edges $P_{tension}^{X,Y}$ for different ratios of junctions with non-zero tension $f_r$ (Fig. S3C,D). The respective percolation thresholds $(f_c) : (\frac{d^2 P_{tension}}{df_r^2} = 0)$ were the inflection points of $P_{tension}$. In Fig. S3B, note that $f_c^x \approx 0.66$ remains unchanged but $f_c^y$ decreases with increasing $\Delta A(t)$. As a result, $f_c = \max[f_c^x, f_c^y]$ remains constant at $f_c \approx 0.66$ throughout dorsal closure. Thus we find that the fraction of junctions with non-zero tension must satisfy $f_r \geq f_c \approx 0.66$ for solid behavior. However, this threshold of 0.66, also found for Voronoi tessellation networks, is only valid for sufficiently large networks (16). For small systems, the probability of obtaining a system-spanning path of junctions with non-zero tension increases dramatically, since shorter paths are needed, and these are more likely than longer paths. This explains the sudden drop of the percolation threshold in the y-direction $f_c^y$ (Fig. 3A in main text).

Uniaxial deformation can also affect $f_r$, the fraction of junctions with non-zero tension. To determine the fraction of junctions with non-zero tension in the model, we chose a threshold corresponding to the noise floor of the tension (17), so that junctions with $\tau_J > 10^{-4}$ are counted as having finite tension. In the experiments, we found that the initial recoil velocity $v_r$ was independent of junction straightness $S$ for $S \lesssim 0.93$ (Fig. 2E, main text), but rose linearly with $S$ above this threshold. We therefore assume that only junctions with $S > 0.93$ carry tension and contribute to solid response. In Fig. S3B it is evident that $f_r > f_c$ throughout the process, consistent with the system being solid.

In summary, the AS tissue maintains a percolating network of tense cell-cell junctions across the dorsal opening during the entire process of closure, consistent with its solid character. The decrease of preferred cell perimeter is crucial. If we leave the preferred perimeter fixed at its initial value in our model, we obtain $f_r$ as shown in Fig. S3B. Clearly, $f_r$ falls below $f_c$ so that the system fluidizes.

**H. Mean junction tension in tissues with and without orientational alignment of the cells.** To understand the effect of the orientational alignment of cells on mechanical stiffness in our model, we compared our model, where we found orientational alignment of cells (Fig. 2C, main text) during uni-axial constriction, with simulations with randomly aligned cells. We then quantified the mean junction tension $\bar{\tau}_J$, as a measure for tissue stiffness, in model configurations with different mean shape indices $\bar{q}$. For a given mean shape index, we find that the junction tension is higher if the cells are oriented than if they are not (Fig. S4), consistent with Ref. (5).

**I. Effect of uniaxial deformation on tissue phase state.** We studied the effect of uniaxial deformation of the AS, as implemented in our model, in contrast to isotropic deformation. We progressively decreased the tissue size in our model in both x and y direction, and compared it with our model results for uniaxial deformation at each given $\Delta A$ in terms of mean shape index $\bar{q}$ (Fig. S5A), average orientational order $\bar{Q}$ (Fig. S5B), average junction tension $\bar{\tau_J}$ and initial recoil velocity $v_J$ (Fig. S5C), and standard deviation of shape index $\sigma_q$ (Fig. S5D). We found that uniaxial deformation of the AS is crucial to recapitulate the experimentally measured time courses of cell-shape features.

To test the effect of uniaxial vs. isotropic deformation on the solid phase of the AS, we assessed the fraction of junctions with nonzero tension $f_r$ during closure (red dashed line in Fig. S3B). For isotropic deformation, our model predicts $f_r$ well above the percolation threshold $f_c$, so non-zero tension junctions percolate in either case (isotropic and uni-axial).

**J. Methodology used to generate the phase diagram.** To construct the phase diagram, we initialized our model for a range of different target shape indices (range from 4.15-4.65, with 15 initial configurations at each target shape indices). We then ran our simulation for these different configurations and each evaluated the mean shape index as well as the fraction of junctions with non-zero tension ($f_r$) at different stages of closure $\Delta A$. This strategy allowed us to sample the phase diagram. In alignment with our own validation and other work, a tissue is solid when $f_r \geq 0.66$. In Figure 3C, we plotted the measured mean shape index against the stage of closure of the individual model states, with open circles representing the solid state and open squares representing the liquid state. The black dashed line represents the phase boundary.

**K. Effect of periodic fluctuations of cell areas on tissue state.** During dorsal closure, amniserosa cells experience oscillations in their apical cell area. We tested how asynchronous periodic fluctuations of preferred cell areas, reminiscent of the pulsatile medioapical contractions of amnioserosa cells, affect tissue-level properties. To test this, we ran a simulation where we introduced a periodic fluctuation in the preferred area term at each point of the closure value, as shown below:

$$\widetilde{a_0[i]} = a_0[i](1 + b\ Sin(wx + \phi)) \qquad [7]$$

where b is the amplitude, x is the simulation step, and a random phase difference is introduced by the $\phi$ term. We used the experimentally measured amplitude, b = 0.09. The total duration of the part of closure we examined is ∼100 minutes, measured experimentally. The period time in the experiment is ∼4 minutes, which corresponds to a total of ∼25 oscillations during the experimental closure process. In the simulation, we have taken total ∼25 oscillations during the closure.

Our results show that the periodic fluctuations only slightly reduced the rigidity of the tissue, but it remained in the rigid phase (Fig. S7A,B). We also evaluated the average shape index during closure and found only a minimal effect (Fig. S7C).

**L. Shear modulus calculation in model tissue.** The shear modulus is measured by evaluating the second derivative of the system's energy with respect to an infinitesimally small applied simple shear strain $\gamma$

$$G = \frac{1}{L^2}\frac{\partial^2 E}{\partial \gamma^2}, \qquad [8]$$

here, $L = \sqrt{N}$ is the dimension of the simulation box. Instead of applying multiple strain deformations and numerically calculating the second derivative, the shear modulus is efficiently obtained from the Hessian matrix associated with the energy-minimized state (18). The elements of the Hessian matrix can be obtained from the second derivatives of the energy (E) with respect to the positions of the vertices.

$$D_{i\alpha,j\beta} = \frac{\partial^2 E}{\partial r_{i\alpha}\partial r_{j\beta}}, \qquad [9]$$

where $r_{i\alpha}$ and $r_{j\beta}$ are the $\alpha$ component of vertex $i$ coordinates and $\beta$ component of vertex $j$ coordinates, respectively. It can be demonstrated that

$$G = \frac{1}{L^2}\left(\frac{\partial^2 E}{\partial \gamma^2} - \sum_m \frac{1}{\omega_m^2}\left[\sum_{j,\alpha}\frac{\partial^2 E}{\partial \gamma \partial r_{j\alpha}} u_{j\alpha}^m\right]^2\right), \qquad [10]$$

where $\omega_m^2$ are the non-zero eigenvalues of the Hessian, $u_{j\alpha}^m$ are the corresponding normalised eigenvectors (18).

**M. Analysis of strain in amnioserosa during dorsal closure.** To quantify the strain accumulated along the x-axis ($\epsilon_{xx}$) and y-axis ($\epsilon_{yy}$) in the amnioserosa during closure, we analyzed experimental data of cell centroid trajectories and neighbor matrices. We followed the procedure described in Ref. (19) (Equations 2.11 to 2.14) to obtain the strain tensor ($\epsilon_{ij}$). Strain components ($\epsilon_{xx}$, $\epsilon_{yy}$, $\epsilon_{xy}$, $\epsilon_{yx}$) were calculated by minimizing the mean-square displacement between the actual displacements of neighboring cells relative to the central cell and the displacements they would have in a region with uniform strain. We computed the isotropic strain (Tr($\epsilon$)) and deviatoric strain ($\sqrt{Tr(\delta\epsilon : \delta\epsilon)}$), where $\delta\epsilon = \epsilon_{ij} - \frac{1}{2}Tr(\epsilon)\delta_{ij}$ (Fig. ??C,D). Fig. S8 presents the strain components $\epsilon_{xx}$ and $\epsilon_{yy}$ as well as isotropic and deviatoric strain as functions of $\Delta A(t)$. The plots show the median strain for individual embryos and the mean across multiple embryos. Note that the initial non-zero offsets of $\epsilon_{xx}$ and $\epsilon_{yy}$ are due to cell centroid fluctuations. Our analysis reveals that $\epsilon_{xx}$ remains nearly constant, indicating only minimal compressive strain along the x-axis. In contrast, $\epsilon_{yy}$ shows a progressive decrease, signifying shrinkage along the y-axis, which aligns with the assumptions in our model. The isotropic and deviatoric strain patterns in our model and experimental data show reasonable agreement.

**N. Analysis of stress in model amnioserosa throughout dorsal closure.** We analyzed the stress tensor ($\sigma_{ij}$) for our model tissue as function of $\Delta A(t)$ following the procedure described in Ref. (20) (Appendix A). We computed the isotropic ($\mathrm{Tr}(\sigma)$) and deviatoric stress ($\sqrt{Tr(\delta\sigma : \delta\sigma)}$), where $\delta\sigma = \sigma_{ij} - \frac{1}{2}Tr(\sigma)\delta_{ij}$. Both components were evaluated as functions of $\Delta A(t)$ as a function of $\Delta A(t)$ (Fig. S9A and B). The persistent non-zero values of both stress components throughout dorsal closure provide evidence for the tissue's solid-like behavior, complementing our percolation studies and shear modulus measurements.

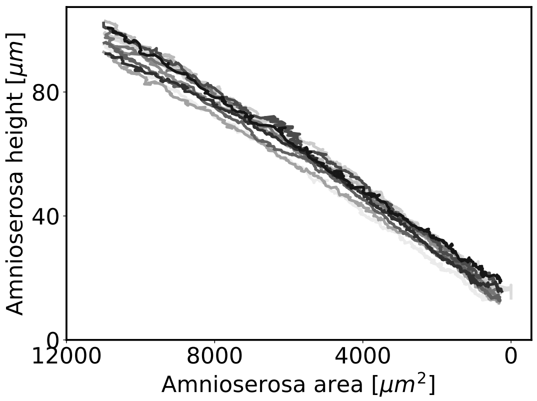


**Fig. S1. Linear relation between the AS height and area.** Each color represents one embryo (N=12).

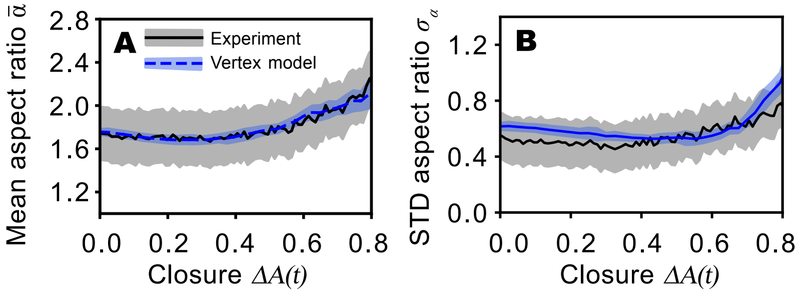


**Fig. S2. Comparison of mean (A) and standard deviation (B) of cell aspect ratio between experimental data (black) and model (blue) during dorsal closure.** Lines show mean, and shaded areas standard deviation of various initial configurations (model, N = 10) or different embryos (experiment, N = 12).

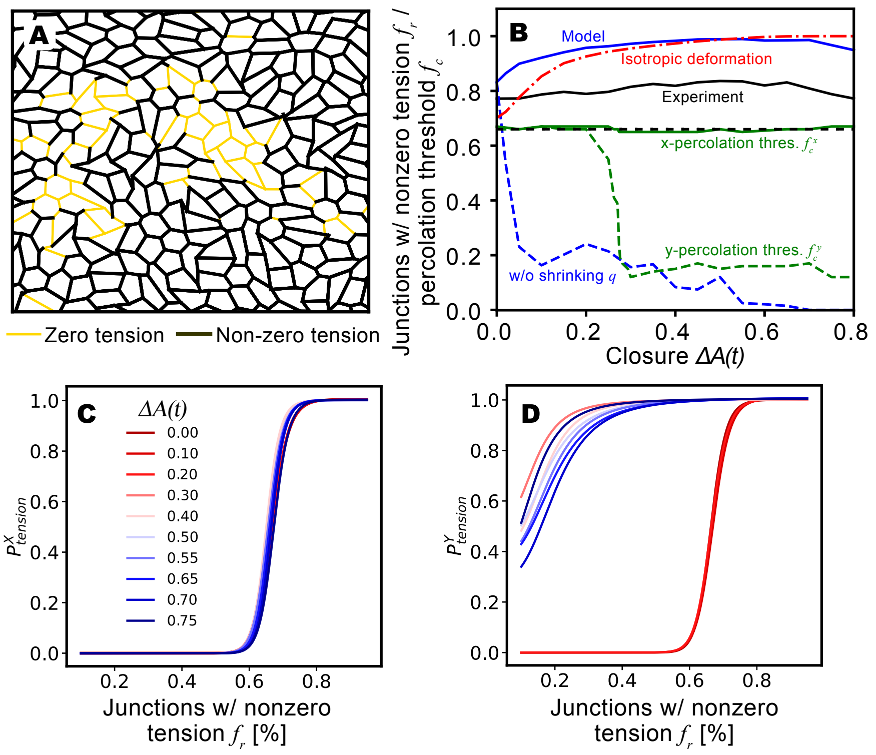


**Fig. S3. Analysis of percolation in junction network.** (**A**) Snapshot of tension network percolation: Black thick lines are edges with finite tensions (percolating throughout the tissue), while other edges have $\tau = 0$. (**B**) Fraction of junctions with nonzero tension, $f_r$, in experiment (black solid) and model (blue solid) as a function of $\Delta A(t)$. The model predicts the minimum fraction $f_c$ needed for percolation of junctions with nonzero tension in the $x$ (green solid) and $y$ (green dashed) directions. In both experiment and model, $f_r > f_c = \max[f_c^x, f_c^y]$, indicating that the system is solid. Black dashed line shows the effective bond percolation threshold $f_c^* = 0.66 = \max[f_c^x, f_c^y]$ above which the system is solid. If cell perimeters, and accordingly the shape indices of cells are held constant instead of shrinking with increasing $\Delta A(t)$, our model predicts that $f_r$ drops below the percolation threshold (blue dashed), leading to fluidization. If we apply isotropic instead of uni-axial deformation, $f_r$ (red dashed) remains above the percolation threshold. (**C**) Probability of obtaining percolation of cell edges with nonzero tensions ($P^X_{tension}$) vs. $f_r$ in the X directions (left) of our model tissue, at various closure stages (legend), which correspond to various tissue anisotropy stages. The point of inflection is the respective percolation threshold $f_c$. (**D**) Same plot as **C** for Y direction, showing substantially lower percolation thresholds than for X direction at late closure stages.

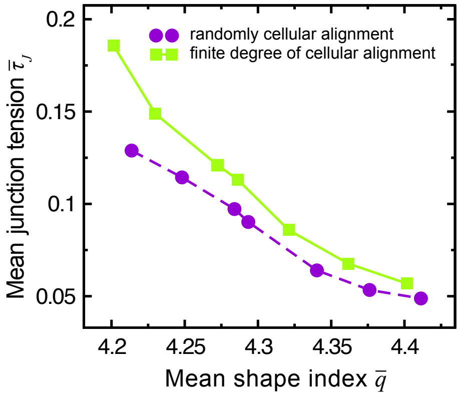


**Fig. S4. Comparison of mean junction tension $\bar{\tau}_J$ for tissues with and without cellular orientational alignment at different mean cell shape indices $\bar{q}$.**

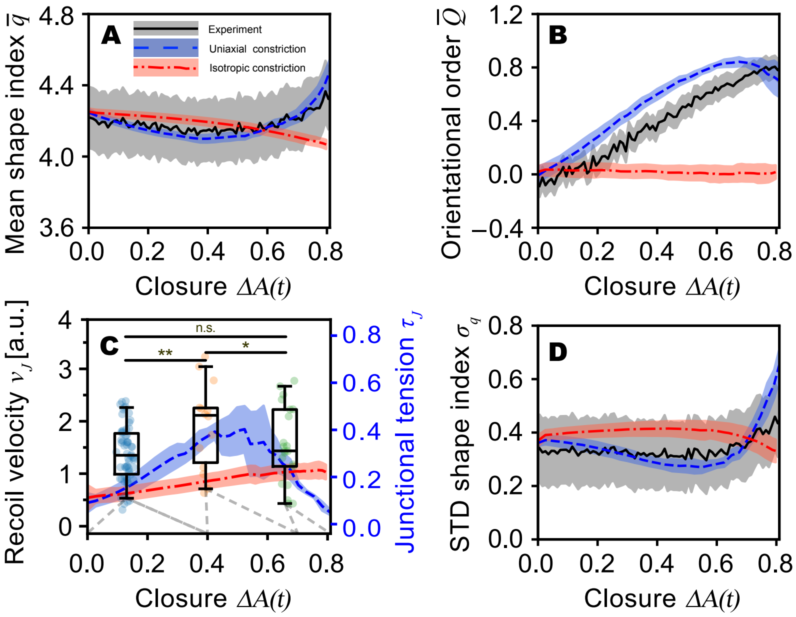


**Fig. S5. Comparison of experimental data (black) with model results with uniaxial deformation (blue) and isotropic deformation (red) during dorsal closure. (A)** Time course of mean shape index $\bar{q}$. **(B)** Time course of average orientational order parameter $\bar{Q}$. **(C)** Box plot of initial junction recoil velocity of junction vertices after performing laser ablation of the junction, and time-course of average junction tension $\bar{\tau}_c$. Boxplots show data for three intervals of $\Delta A$ ($\Delta A < 0.4$, $0.4 \leq \Delta A < 0.7$, $\Delta A \geq 0.7$). Whiskers extend to the 5th and 95th percentiles, while the boxes delineate the interquartile range, and the horizontal lines within the boxes indicate the median values. An ANOVA followed by a post-hoc Tukey's HSD test was conducted to assess statistical significance (*: $p < 0.1$, **: $p < 0.05$). We performed and evaluated cuts of $N = 97$ junctions. **(D)** Time-course of standard deviation of shape index ($\sigma_q$). Lines show mean, and equally color intervals show respective standard deviations (N = 10 model initial configurations, N = 12 embryos).

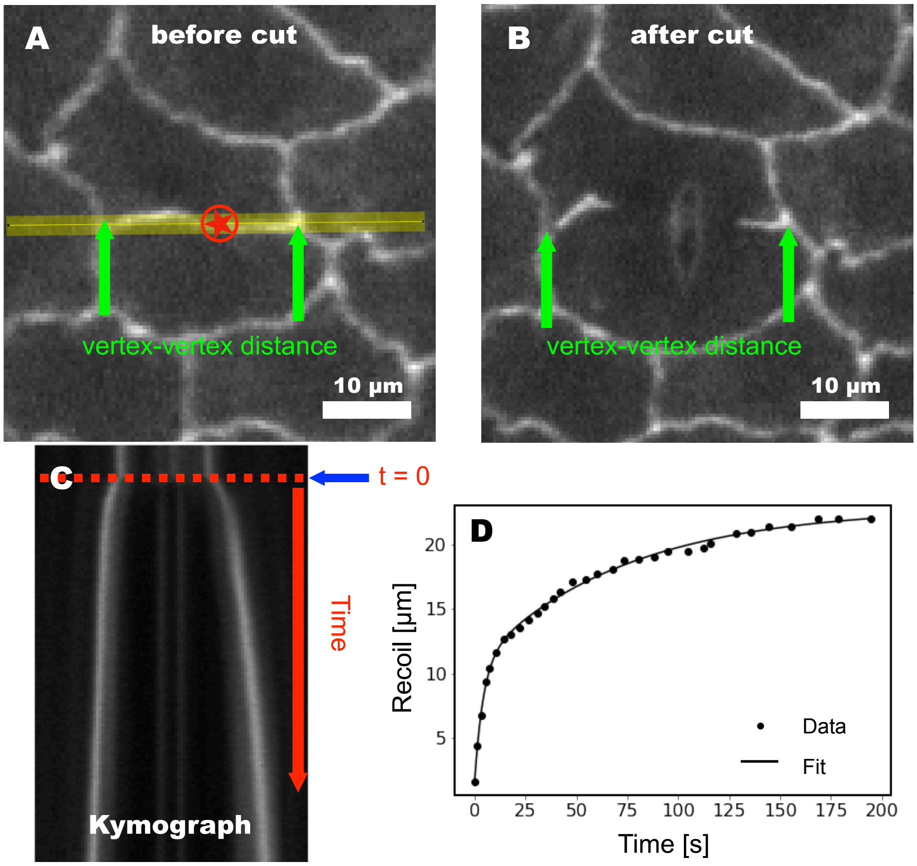


**Fig. S6. Measurement and evaluation of initial recoil velocity of vertices after laser cut.** (**A**),(**B**) Confocal fluorescent images of AS cells. The red star in (A) marks the ablation region. Green arrows show the vertex positions just prior to (A) and after (B) ablation. The yellow line (A) along which the motion of vertices is assessed using kymograph analysis. (**C**) Kymograph of positions of the two vertices showing recoil after ablation (the red dotted line shows time of cut at $t = 0$). (**D**) Time-course of recoil after ablation. The recoil was well fitted by a double exponential function $a_0 \exp(b_0 t) + c_0 \exp(d_0 t) + e_0$.

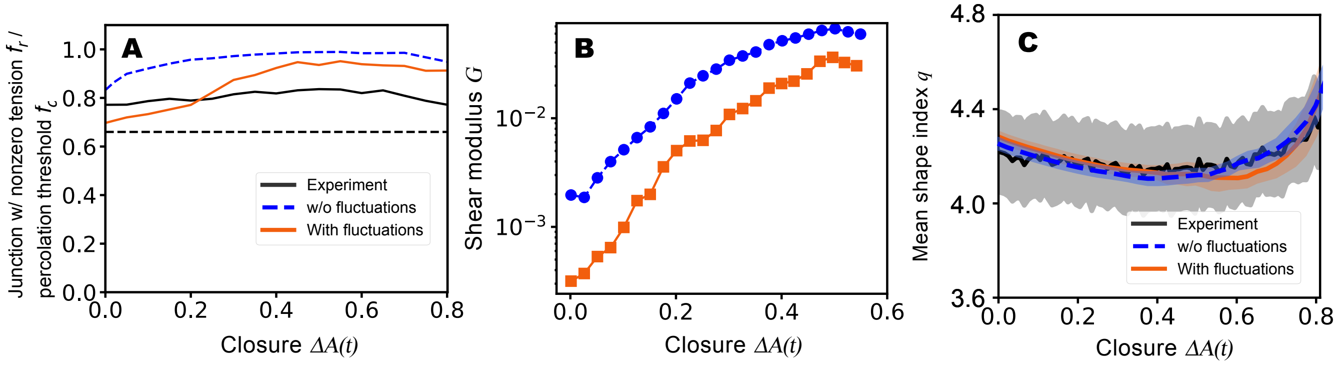


**Fig. S7. Effect of cell area fluctuations on tissue rigidity and cell shapes (A)** Fraction of junctions with nonzero tension $f_r$, in experiment (black solid) and model with periodic fluctuations of preferred cell area (red solid) and without periodic fluctuations of preferred cell area (blue dashed) as a function of $\Delta A(t)$. The Black dashed line shows the effective bond percolation threshold $f_c^*$ = 0.66 above which the system is solid. **(B)** Shear modulus (G) as a function of $\Delta A(t)$ with periodic fluctuations (red solid) and without periodic fluctuations (blue dashed). **(C)** Comparison of shape index with and without periodic fluctuations of preferred cell area as a function of $\Delta A(t)$.

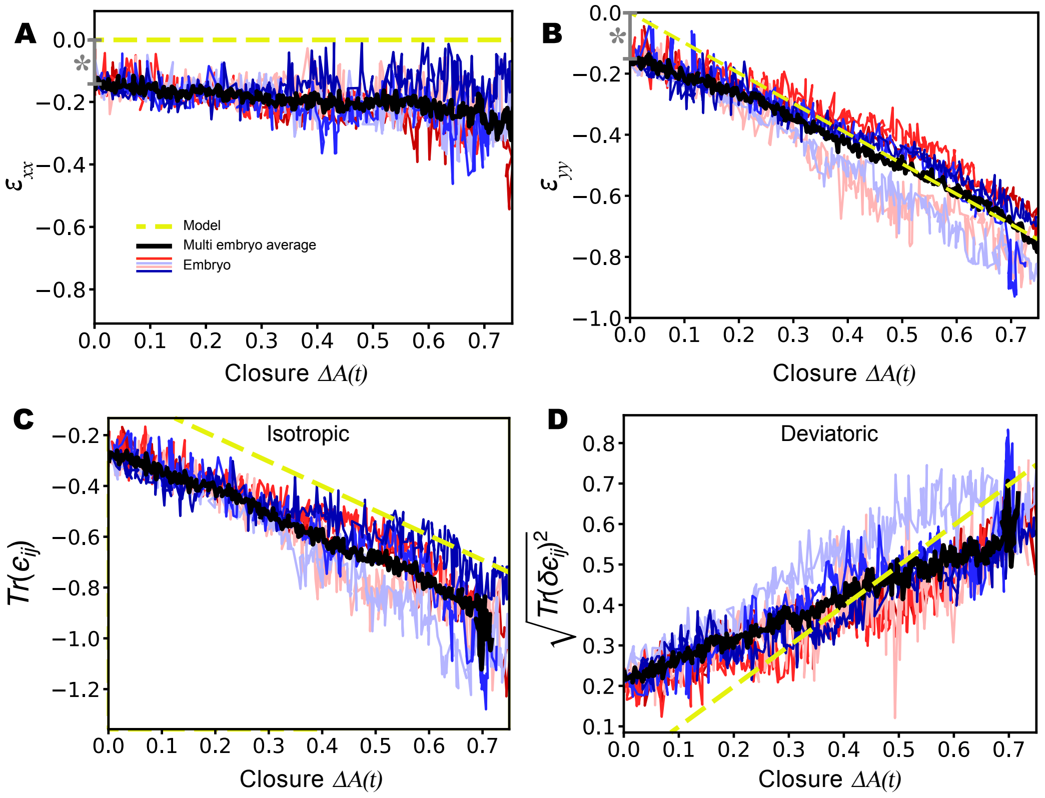


**Fig. S8. Analysis of amnioserosa strain during closure**. In all panels, black lines represent the average across embryos, thin colored lines show individual embryos, and yellow dashed lines indicate the model prediction. **(A)** Strain tensor along x-axis ($\epsilon_{xx}$) as a function of $\Delta A(t)$. The asterisk marks an offset due to cell centroid fluctuations, an artifact of the centroid-based method used to calculate strain. **(B)** Strain tensor along y-axis ($\epsilon_{yy}$) over time. The model curve shows the anisotropic shrinkage assumed, closely matching experimental observations. The asterisk indicates the offset due to cell centroid fluctuations as in (A). **(C)** Isotropic strain over time. **(D)** Deviatoric strain over time. Data shown are from N=6 embryos, in which the horizontal image axis was well aligned with the anterior-posterior axis.

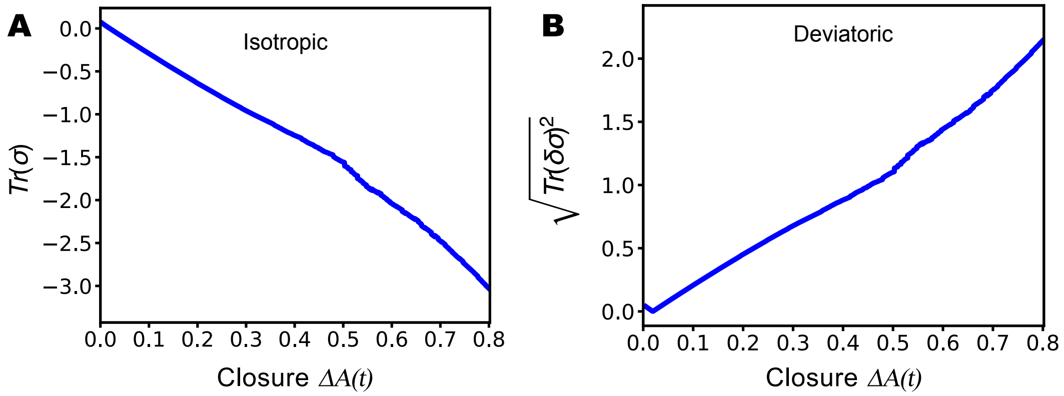


**Fig. S9. Analysis of isotropic and deviatoric stress in model tissue(A)** The isotropic stress as a function of $\Delta A(t)$. **(B)** The deviatoric stress as a function of $\Delta A(t)$.

### Legends for Movies

**Movie S1.** Time-lapse confocal microscopy of a dorsal closure stage embryo expressing GFP tagged E-cadherin to outline the amniosersosa cell boundaries.. Yellow dots show centroids of cells included in model-experiment comparison. Scale bar represents 50 µm. The frame rate is 4 frames/minute.
